# *Neotrygon vali*, a new species of the blue-spotted maskray complex (Myliobatoidei: Dasyatidae)

**DOI:** 10.1101/106682

**Authors:** Philippe Borsa

## Abstract

The blue-spotted maskray from Guadalcanal Island (Solomon archipelago) is distinct by its colour patterns from *Neotrygon kuhlii* with which it was previously confused, and belongs to a genetic lineage clearly separate from all other known species in the genus *Neotrygon*. It is here described as a new species, *Neotrygon vali* sp. nov., on the basis of its nucleotide sequence at the *cytochrome oxidase 1* (*CO1*) gene locus. It is diagnosed from all other known species in the genus *Neotrygon* by the possession of nucleotide T at nucleotide site 420 and nucleotide G at nucleotide site 522 of the *CO1* gene.

## INTRODUCTION

Genetic studies of the dasyatid genus *Neotrygon* Castelnau, 1873 or maskrays have pointed to the possible occurrence of several species complexes (Ward et al., 2008; Naylor et al., 2012; Borsa et al., 2016a and references therein). This genus currently comprises 10 nominal species: *N. annotata* (Last, 1987), *N. australiae* Last, White and Séret, 2016, *N. caeruleopunctata* Last, White and Séret, 2016, *N. kuhlii* (Müller and Henle, 1841), *N. leylandi* (Last, 1987), *N. ningalooensis* Last, White and Puckridge, 2010, *N. orientale* Last, White and Séret, 2016, *N. picta* (Last, 1987), *N. trigonoides* (Castelnau, 1873) and *N. varidens* (Garman, 1885). The bluespotted maskray, previously *N. kuhlii*, consists of up to eleven lineages representing separate species (Arlyza et al., 2013a; Puckridge et al., 2013; Borsa et al., 2016a, 2016b) of which four (*N. australiae*, *N. caeruleopunctata*, *N. orientale*, *N. varidens*) have so far been formally described. One of the paratypes of *N. kuhlii*, a specimen from Vanikoro in the Santa Cruz archipelago, has been recently designated as lectotype (Last et al., 2016), although the pigmentation patterns of the Vanikoro maskray, thus now the typical *N. kuhlii*, do not fit those of the original description of the species by J. Müller and F.G.J. Henle (Müller and Henle, 1841; Borsa and Béarez, 2016). In their re-description of *N. kuhlii*, Last et al. (2016) hastily included a fresh specimen collected from Guadalcanal Island in the Solomon archipelago, over 800 km away from Vanikoro, the type-locality. Pigmentation patterns clearly distinguish the Guadalcanal maskray from *N. kuhlii* from Vanikoro (Borsa and Béarez 2016), but not from other species previously under *N. kuhlii* except *N. varidens* (Garman 1885).

In contrast, mitochondrial DNA sequence information contributes valuable diagnostic characters to the taxonomic description of species and is fundamental to the description of cryptic species (Jörger and Schrödl, 2013). The taxonomic value of mitochondrial DNA sequences has been demonstrated in morphologically intractable species complexes in Elasmobranchs such as *Himantura uarnak* and *N. kuhlii* (Naylor et al., 2012; Arlyza et al., 2013a; Borsa et al., 2013a, 2013b; Puckridge et al., 2013; Borsa, 2017).

It is here emphasized that after careful re-examination of Last et al.’s (2016) work, Borsa et al. (in press) found no diagnostic morphological character that clearly distinguished any of the three new species described from the two others or from *N. kuhlii*. Thus, Last et al.’s (2016) morphological diagnoses were found to be invalid. The objectives of the present paper, which follows up Borsa and Béarez (2016), are the following: (1) to identify diagnostic characters that distinguish the Guadalcanal maskray from other species in the genus *Neotrygon*; (2) to describe it as a new maskray species, a necessary step towards clarifying the intricate taxonomy of species in this species complex.

## METHODS

Because *N. kuhlii* from Vanikoro, the type-locality, has not yet been analyzed genetically, pigmentation patterns were used to distinguish it from the Guadalcanal maskray, following Borsa et al. (2013a). Three specimens of the Guadalcanal maskray were examined including specimen no. CSIRO H 7723-01 (p. 539 of Last et al., 2016) and two live specimens photographed underwater, one by Randall (2005) and the other one by M.A. Rosenstein (Fig. 1). The diameter of ocellated blue spots on the dorsal side of the disk, relative to disk width, was measured on the photographs. Ocellated blue spots were qualified as “small” when their maximum diameter was ≤ 2% disk width (DW), “medium” when ≤ 4% DW and “large” when > 4% DW (Borsa et al., 2013a). On Randall’s (2005) picture and on Fig. 1, DW was deduced from disk length (DL; measured from tip of snout to rear tip of pelvic fin) from the relationship DW = 1.13 DL, obtained from measurements on specimen no. CSIRO H 7723-01. Dark speckles (≤ 1% DW) and dark spots (> 1% DW) were also counted on the dorsal surface of the disk (Borsa et al., 2013a). The counts did not include those speckles and spots located within the dark band around eyes that forms the mask. The presence or absence of a scapular blotch was also checked.

**Figure 1.**
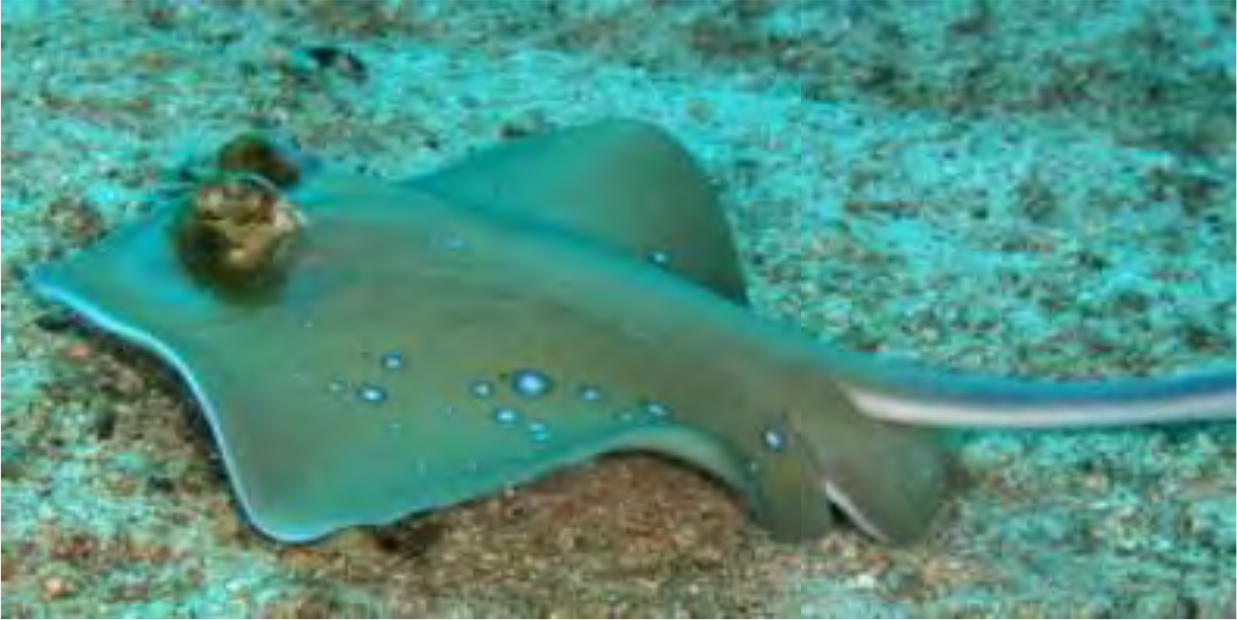
Guadalcanal maskray *Neotrygon vali* sp. nov. showing the pigmentation patterns that differentiate it from *N. kuhlii* from Vanikoro (Borsa and Béarez 2016). Photographed by M.A. Rosenstein near Mbike Wreck (09°06’S 160°11E), November 2014.

The Guadalcanal maskray was compared to other species in the genus *Neotrygon* based on nucleotide sequences of the *CO1* gene. A total of 205 complete or partial *CO1* gene sequences were found in the literature (Ward et al., 2008; Yagishita et al., 2009; Aschliman et al., 2012; Arlyza et al., 2013a; Borsa et al., 2013a; Puckridge et al., 2013; Last et al., 2016) and compiled into a single FASTA file which was edited under BIOEDIT (Hall, 1999). The recently-described *N. australiae* and *N. caeruleopunctata* correspond to, respectively, clades *V* and *VI* of Arlyza et al. (2013a). Clade *IV* of Arlyza et al. (2013a) included a distinct sub-clade that corresponds to *N. varidens*. All other haplotypes of clade *IV* of Arlyza et al. (2013a), together with GenBank no. JN184065 (Aschliman et al., 2012) correspond to *N. orientale*, except a distinct haplotype (GenBank no. AB485685; Yagishita et al., 2009) here referred to as the Ryukyu maskray. Two haplotypes from the Indian Ocean (GenBank nos. JX263421 and KC249906) belonging to Haplogroup *I* of Arlyza et al. (2013a) are here referred to as the Indian Ocean maskray. Sample sizes were: *N* = 8 for *N. annotata*; *N* = 11 for *N. australiae*; *N* = 12 for *N. caeruleopunctata*; *N* = 7 for *N. leylandi*; *N* = 1 for *N. ningalooensis*; *N* = 68 for *N. orientale*; *N* = 5 for *N. picta*; *N* = 18 for *N. trigonoides*; *N* = 11 for *N. varidens*; *N* = 19 for clade *II* of Arlyza et al. (2013a); *N* = 17 for clade *III* of Arlyza et al. (2013a); *N* = 14 for clade *VII* of Arlyza et al. (2013a); *N* = 10 for clade *VIII* of Arlyza et al. (2013a); *N* = 1 for the Guadalcanal maskray; *N* = 2 for the Indian Ocean maskray; and *N* = 1 for the Ryukyu maskray. GenBank accession numbers for all the foregoing sequences are provided in Supplementary Table S1.

Average nucleotide divergences between pairs of sequences within a lineage and net nucleotide divergences between lineages were estimated according to the Tamura-3 parameter substitution model (Tamura, 1992), the most likely model as inferred from the Bayesian information criterion using MEGA6 (Tamura et al., 2013). Variable nucleotide sites were determined automatically using MEGA6. Diagnostic nucleotide sites at the *CO1* gene locus that distinguish the Guadalcanal maskray from all other lineages in the genus *Neotrygon* were then selected visually on the EXCEL (Microsoft Corporation, Redmond WA) file generated by MEGA6.

## RESULTS AND DISCUSSION

Last et al. (2016) have claimed that the Guadalcanal maskray specimen they had in hands was “very similar in coloration and shape to Müller and Henle’s Solomon Island types” but this statement was shown to be unwarranted (Borsa and Béarez, 2016). Pigmentation patterns on the dorsal side of each pectoral fin in the Guadalcanal maskray consisted of a variable number (*N* = 2-21) of small ocellated blue spots, a small number (*N* = 1-6) of medium-sized ocellated blue spots, and 3-7 dark speckles (Table 1). All three Guadalcanal maskray specimens available for the present study thus lacked the dark spots and the scapular blotch that are present in the Vanikoro maskray, i.e. *N. kuhlii* (Borsa and Béarez, 2016). Given the relevance of pigmentation patterns in diagnosing species in the genus *Neotrygon* (Last and White, 2008; Last et al., 2010; Borsa et al., 2013a) and more generally in stingrays (Arlyza et al., 2013b; Borsa, 2017), this observation alone suffices to reject the hypothesis that the Guadalcanal maskray is synonymous with *N. kuhlii*. Other measurements, expressed as percentage of disc length (DL), also showed strong differences between the Guadalcanal maskray and the type material of *N. kuhlii* including the lectotype (MNHN-IC-0000-2440, smaller of two) and the paralectotype (MNHN-IC-0000-2440, larger of two). For instance, the distance from pectoral fin insertion to sting origin was substantially larger in the Guadalcanal maskray (5.4% DL) than in *N. kuhlii* (4.2% DL), as was the nostril length (5.0% DL *vs*. 3.4-3.9% DL). The inter-orbital width was substantially narrower (9.2% DL *vs*. 10.3-11.6% DL), as were the inter-ocular width (19.7% DL *vs*. 21.3-22.6% DL), the distance between first-gill slits (19.2% DL *vs*. 21.9% DL), and the distance between fifth-gill slits (9.8% DL *vs*. 11.1% DL).

**Table 1.**
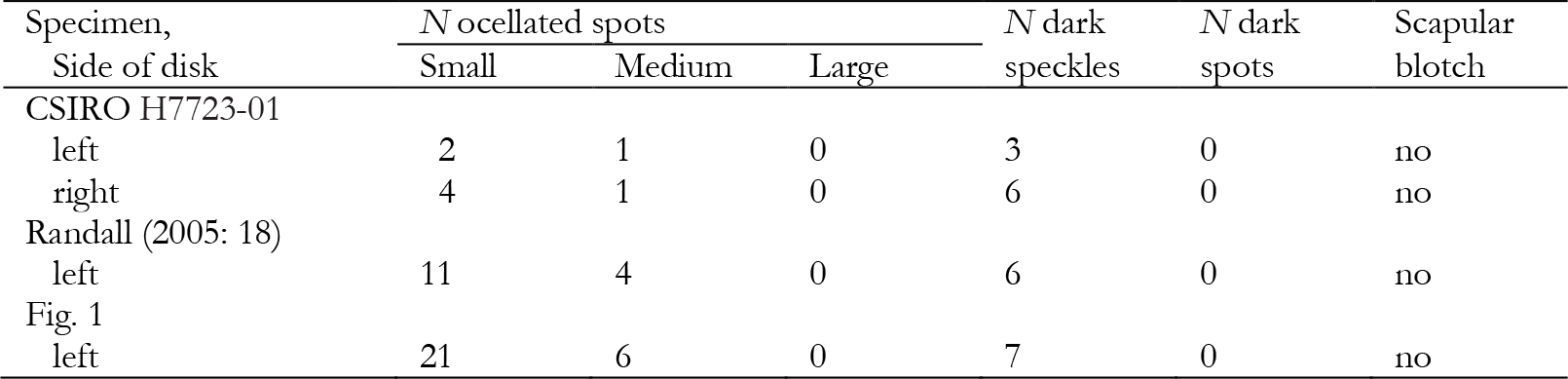
Pigmentation patterns on left or right dorsal side of disk in Guadalcanal maskray *Neotrygon vali* sp. nov. including numbers of ocellated blue spots, number of dark speckles or spots and presence or absence of a scapular blotch. Ocellated blue spots qualified as *small* when diameter ≤ 2% disk width (DW); *medium* when > 2% DW and ≤ 4% DW and *large* when > 4% DW; *dark speckles* ≤ 1% DW; *dark spots* > 1% DW (Borsa et al., 2013a). *N*: number of speckles or spots.

| Specimen,<br>Side of disk | *N* ocellated spots |  |  | *N* dark<br>speckles | *N* dark<br>spots | Scapular<br>blotch |
| --- | --- | --- | --- | --- | --- | --- |
|  | Small | Medium | Large |  |  |  |
| CSIRO H7723-01 |  |  |  |  |  |  |
| left | 2 | 1 | 0 | 3 | 0 | no |
| right | 4 | 1 | 0 | 6 | 0 | no |
| Randall (2005: 18) |  |  |  |  |  |  |
| left | 11 | 4 | 0 | 6 | 0 | no |
| Fig. 1 |  |  |  |  |  |  |
| left | 21 | 6 | 0 | 7 | 0 | no |

The maximum-likelihood tree of *CO1* haplotypes (Fig. 2) confirmed the monophyly of species in the genus *Neotrygon*, except *N. picta* which was paraphyletic with *N. leylandi*. Also, no distinction was evident between haplotypes of *N. annotata* and those previously assigned to a related undescribed lineage provisionally referred to as “*Neotrygon* cf. *annotata*” (Puckridge et al., 2013). Estimates of nucleotide divergence at the *CO1* locus among species and deep lineages [i.e. cryptic species remaining undescribed; Borsa et al. (2016b)] in the genus *Neotrygon* ranged from 0.015 to 0.301 (Table 2). They ranged from 0.015 to 0.038 among the four already-described blue-spotted maskray species previously under *N. kuhlii*, i.e. *N. australiae*, *N. caeruleopunctata*, *N. orientale* and *N. varidens* (Table 2). Nucleotide divergence between the Guadalcanal maskray and other species in the genus *Neotrygon* was ≥ 0.049 (Table 2). Meanwhile, nucleotide divergence estimates within lineages ranged from 0 in *N. caeruleopunctata* to 0.011 in *N. orientale* and in clade *II* of Arlyza et al. (2013a) (Table 2), thus systematically lower than inter-specific estimates, and largely so. The single Guadalcanal maskray haplotype belonged to a lineage clearly distinct from all other *Neotrygon* spp. lineages sampled so far. At two sites at the *CO1* locus, it possessed nucleotides that were absent in *N. annotata*, *N. australiae*, *N. caeruleopunctata*, *N. leylandi*, *N. ningalooensis*, *N. orientale*, *N. picta*, *N. trigonoides*, *N. varidens*, and in six yet-undescribed blue-spotted maskray species sampled from the Indian Ocean, the western and northern costs of Sumatra, the Malacca strait, the Banda sea, the Ryukyu archipelago and West Papua (Arlyza et al., 2013a; Borsa et al., 2016a, 2016b) (Supplementary Table S1). Nucleotide sequences at the *CO1* locus therefore provided diagnostic characters for the Guadalcanal maskray, relative to all other species in the genus *Neotrygon*. The Guadalcanal maskray is here considered to represent a distinct species, based on its colour patterns, its distinct phylogenetic placement, its level of nucleotide distance with other species in the genus *Neotrygon*, and its unique nucleotide composition at the *CO1* locus. No name being available for the Guadalcanal maskray (Eschmeyer et al., 2016), it is here described as a new species.

**Figure 2.**
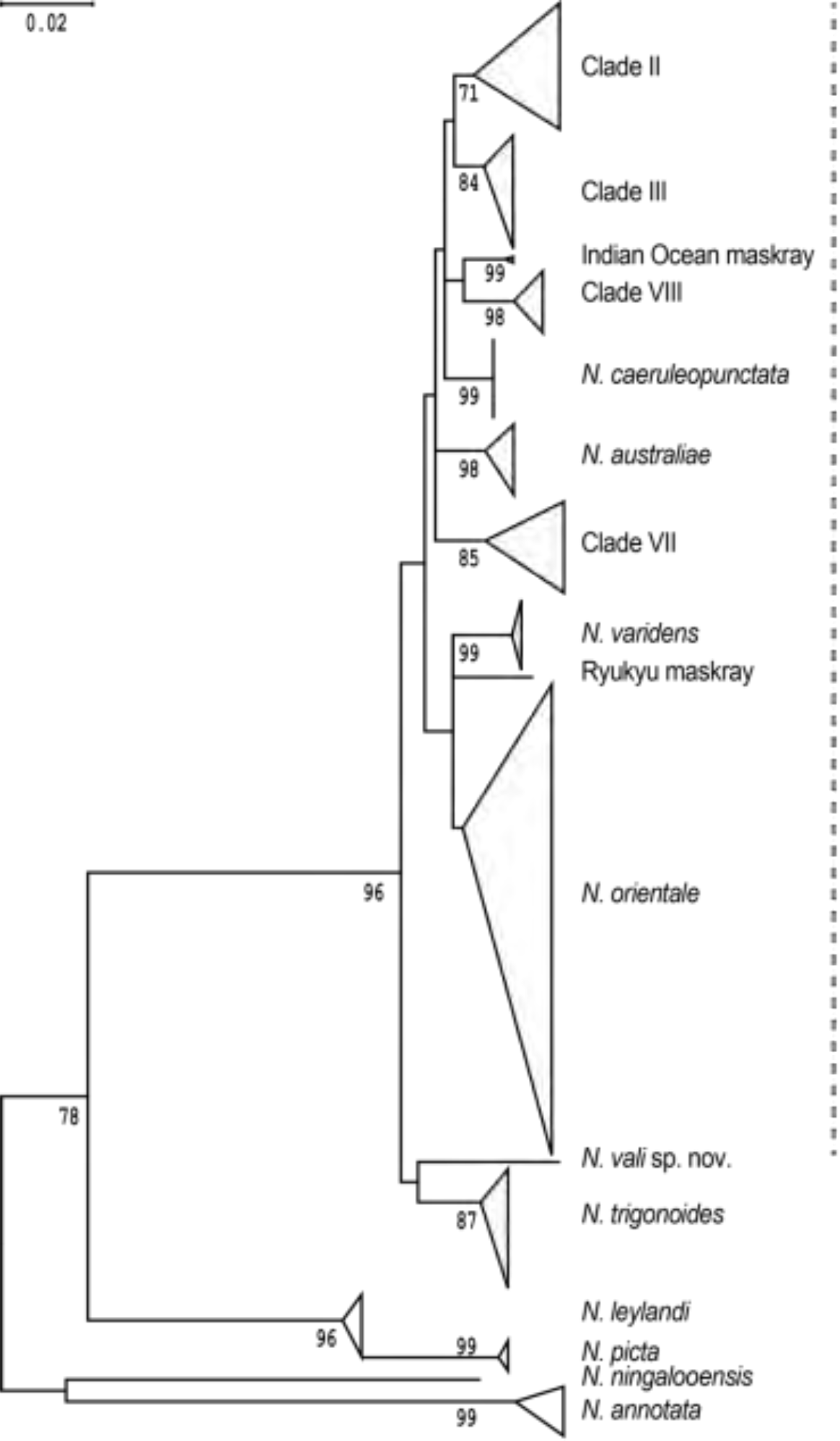
*Neotrygon* spp. Maximum-likelihood tree (Tamura 3-parameter model; MEGA6) of nucleotide sequences at the *CO1* locus (*N* = 205), compiled from several sources (Ward et al., 2008; Yagishita et al., 2009; Arlyza et al., 2013a; Borsa et al., 2013a; Puckridge et al., 2013; Last et al., 2016; Aschliman et al., 2012) showing the phylogenetic placement of the Guadalcanal maskray *Neotrygon vali* sp. nov. Numbers at nodes are bootstrap scores (500 bootstrap resampling runs under MEGA6). Dotted vertical line: blue-spotted maskrays previously under *N. kuhlii* (Borsa et al., 2016b).

**Table 2.** *Neotrygon* spp. Estimates of net nucleotide divergence (Tamura-3 parameter model; MEGA6) between lineages. Clades *II*, *III*, *VII* and *VIII* were defined by Arlyza et al. (2013a). *N* sample size; *ns* number of base substitutions per site from averaging over all sequence pairs within each lineage (Tamura-3 parameter model; MEGA6).

| No. | Lineage | *N* | *ns* | Lineage no. |  |  |  |  |  |  |  |  |  |  |  |  |  |  |
| --- | --- | --- | --- | --- | --- | --- | --- | --- | --- | --- | --- | --- | --- | --- | --- | --- | --- | --- |
|  |  |  |  | 1 | 2 | 3 | 4 | 5 | 6 | 7 | 8 | 9 | 10 | 11 | 12 | 13 | 14 | 15 |
| 1 | *N. annotata* | 8 | 0.004 |  |  |  |  |  |  |  |  |  |  |  |  |  |  |  |
| 2 | *N. australiae* | 11 | 0.006 | 0.268 |  |  |  |  |  |  |  |  |  |  |  |  |  |  |
| 3 | *N. caeruleopunctata* | 12 | 0.000 | 0.278 | 0.028 |  |  |  |  |  |  |  |  |  |  |  |  |  |
| 4 | *N. leylandi* | 7 | 0.002 | 0.243 | 0.167 | 0.179 |  |  |  |  |  |  |  |  |  |  |  |  |
| 5 | *N. ningalooensis* | 1 | - | 0.229 | 0.271 | 0.267 | 0.201 |  |  |  |  |  |  |  |  |  |  |  |
| 6 | *N. orientale* | 68 | 0.011 | 0.236 | 0.029 | 0.028 | 0.160 | 0.233 |  |  |  |  |  |  |  |  |  |  |
| 7 | *N. picta* | 5 | 0.001 | 0.286 | 0.190 | 0.205 | 0.034 | 0.213 | 0.174 |  |  |  |  |  |  |  |  |  |
| 8 | *N. trigonoides* | 18 | 0.003 | 0.250 | 0.047 | 0.044 | 0.178 | 0.212 | 0.036 | 0.174 |  |  |  |  |  |  |  |  |
| 9 | *N. vali* sp. nov. | 1 | - | 0.301 | 0.054 | 0.049 | 0.192 | 0.235 | 0.053 | 0.193 | 0.054 |  |  |  |  |  |  |  |
| 10 | *N. varidens* | 11 | 0.001 | 0.240 | 0.036 | 0.038 | 0.161 | 0.269 | 0.015 | 0.189 | 0.047 | 0.064 |  |  |  |  |  |  |
| 11 | Clade *II* | 19 | 0.011 | 0.288 | 0.027 | 0.021 | 0.199 | 0.263 | 0.029 | 0.214 | 0.035 | 0.034 | 0.050 |  |  |  |  |  |
| 12 | Clade *III* | 17 | 0.003 | 0.282 | 0.027 | 0.021 | 0.198 | 0.266 | 0.024 | 0.203 | 0.044 | 0.038 | 0.016 | 0.043 |  |  |  |  |
| 13 | Clade *VII* | 14 | 0.008 | 0.262 | 0.028 | 0.027 | 0.154 | 0.220 | 0.028 | 0.194 | 0.039 | 0.039 | 0.027 | 0.027 | 0.050 |  |  |  |
| 14 | Clade *VIII* | 10 | 0.002 | 0.249 | 0.028 | 0.022 | 0.150 | 0.251 | 0.034 | 0.175 | 0.037 | 0.044 | 0.025 | 0.027 | 0.025 | 0.056 |  |  |
| 15 | Indian O. maskray | 2 | 0.002 | 0.271 | 0.031 | 0.026 | 0.169 | 0.245 | 0.031 | 0.201 | 0.052 | 0.043 | 0.024 | 0.025 | 0.018 | 0.020 | 0.049 |  |
| 16 | Ryukyu maskray | 1 | - | 0.246 | 0.039 | 0.038 | 0.173 | 0.250 | 0.024 | 0.187 | 0.039 | 0.032 | 0.028 | 0.038 | 0.039 | 0.038 | 0.049 | 0.041 |

## TAXONOMY

Maskrays, genus *Neotrygon* Castelnau, 1873 belong to family Dasyatidae Jordan, 1888. The type species of the genus is *N. trigonoides* (Castelnau, 1873) previously resurrected from synonymy with *N. kuhlii* (Borsa et al., 2013a).

*Neotrygon vali* sp. nov. http://zoobank.org/A5BE7B5D-64A3-40C2-AD44-63ECAE060FF6. Previously referred to as: Guadalcanal maskray (Borsa and Béarez, 2016; Borsa et al., 2016b; Borsa et al., in press); erroneously placed under *Neotrygon kuhlii* by Last et al. (2016).

*Holotype*. Specimen CSIRO H 7723-01, a female 295 mm DW, is here designated as the holotype of *Neotrygon vali* sp. nov. This specimen was obtained on 7 May 2015 from the Plaza fish market, Honiara, Guadalcanal Island (Last et al., 2016). Based on the assumption that fishes sold at the local fish market in Honiara have been captured along the shores of Guadalcanal Island, the type locality is Guadalcanal Island in the Solomon archipelago.

*Description*. The morphological description of the holotype of *Neotrygon vali* sp. nov. has been published previously (pp. 535-541 of Last et al., 2016). This includes 11 meristic counts and 40 measurements made on the body (table 1 of Last et al., 2016). In addition, pigmentation patterns on the dorsal side of disk consist of a variable number of small ocellated blue spots and a moderate number of medium-sized ocellated blue spots, few dark speckles and no scapular blotch. The *CO1* gene sequence of *Neotrygon vali* sp. nov. is unique among species in the genus *Neotrygon* as it clusters with no one of its homologues in congeneric species (Fig. 2). The partial *CO1* gene sequence of the holotype, comprised between homologous nucleotide sites nos. 95 and 696 of the *CO1* gene in *N. orientale* (GenBank no. JN184065; Aschliman et al., 2012) is 5’-C T G G C C T C A G T T T A C T T A T C C G A A C A G A A C T A A G C C A A C C A G G C G C T T T A C T G G G T G A T G A T C A G A T T T A T A A T G T A A T C G T T A C T G C C C A C G C C T T C G T A A T A A T C T T C T T T A T A G T A A T A C C A A T T A T A A T C G G T G G G T T T G G T A A C T G A C T A G T G C C C C T G A T G A T T G G A G C T C C G G A C A T A G C C T T T C C A C G A A T A A A C A A C A T A A G T T T C T G A C T T C T G C C T C C C T C C T T C C T A T T A C T G C T A G C C T C A G C A G G A G T A G A A G C C G G A G C C G G A A C A G G T T G A A C A G T T T A T C C T C C A T T A G C T G G T A A T C T A G C A C A T G C T G G A G C T T C T G T G G A C C T T A C A A T C T T C T C T C T T C A C C T A G C A G G T G T T T C C T C T A T T C T G G C A T C C A T C A A C T T T A T C A C A A C A A T T A T T A A T A T A A A A C C G C C T G C A A T C T C C C A A T A T C A A A C C C C A T T A T T C G T C T G A T C C A T C C T T G T T A C A A C T G T G C T T C T C C T G C T A T C C C T A C C A G T C C T A G C A G C T G G C A T T A C T A T A C T C C T C A C A G A C C G A A A T C T T A A T A C A A C T T T C T T T G A T C C A G C T G G A G G A G G A G A T C C T A T T C T T T A C −3’ (Last et al., 2016).

*Diagnosis*. Based on Supplementary Table S1, *Neotrygon vali* sp. nov. is distinguished from all other species in the genus *Neotrygon* except *N. kuhlii* for which no genetic information is available yet, by the possession of nucleotide T at nucleotide site 420 and G at nucleotide site 522 of the *CO1* gene. In addition, the Guadalcanal maskray is distinct from *N. kuhlii* by the lack of dark spots (> 1% DW) and by the lack of a pair of scapular blotches on the dorsal side.

*Distribution*. Apart from the type locality (Honiara on the northern coast of Guadalcanal Island in the Solomon archipelago), the distribution of *Neotrygon vali* sp. nov. is likely to be confined within the part of Melanesia east of Cenderawasih Bay in West Papua, where the lineage present is *Neotrygon* clade *VIII* (Arlyza et al., 2013a) and west of the Santa Cruz archipelago, where the species present is *N. kuhlii*.

*Etymology. “*Vali” is the word for stingray in Gela, one of the languages spoken in Guadalcanal (Froese and Pauly, 2016). Epithet *vali* is intended to refer to the common name of the species among Guadalcanal fishers and it is a noun in apposition (Truper and De’Clari, 1997). Proposed vernacular names: Guadalcanal maskray (English); vali Guadalcanal (Gela); pastenague masquée à points bleus de Guadalcanal (French).

*Notice*. The present article in portable document (.pdf) format is a published work in the sense of the International Code of Zoological Nomenclature (International Commission on Zoological Nomenclature 1999) or Code and hence the new names contained herein are effectively published under the Code. This published work and the nomenclatural acts it contains have been registered in ZooBank (http://zoobank.org/), the online registration system for the International Commission on Zoological

Nomenclature. The ZooBank life science identifier (LSID) for this publication is urn:lsid:zoobank.org:pub:69E3F1C8-1137-4EF9-B61A-5B56667477A3. The online version of this work is archived and available from the *bioRxiv* (http://biorxiv.org/) and *haL-IRD* (http://www.hal.ird.fr/) repositories.

## CONFLICTS OF INTEREST

No one.

## ACKNOWLEDGEMENTS

I am grateful to P. Béarez (MNHN, Paris), N. Hubert (IRD, Cibinong) and B. Ward (CSIRO, Hobart) for stimulating discussions; to Y. Yates (Tulagi Dive, Honiara) for helpful information; and to M. Rosenstein (ActWin, Cambridge MA) for kindly allowing me to use his underwater photograph of Guadalcanal maskray. I am also grateful to P. Béarez and L. Randrihasipara for high-definition photographs of the Vanikoro syntypes of *Trygon kuhlii*. Insightful comments from four anonymous reviewers were appreciated (Supplementary Table S2). Libel (see Last review of Supplementary Table S2) was taken as encouragement to persevere. Nineteenth-century books and articles were consulted online from the Biodiversity Heritage Library website (http://www.biodiversitylibrary.org/). Authors’ manuscript versions of a series of previous papers on the genetics and taxonomy of the blue-spotted maskray species complex are available from the open-access haL-IRD website (http://www.hal.ird.fr/). This is a contribution of the PARI project, a cooperative research project by IRD, France and LIPI, Indonesia. I declare no conflict of interest and no specific funding for the writing of this paper, of which I am entirely responsible.

## Appendices

**Supplementary Table S1.**
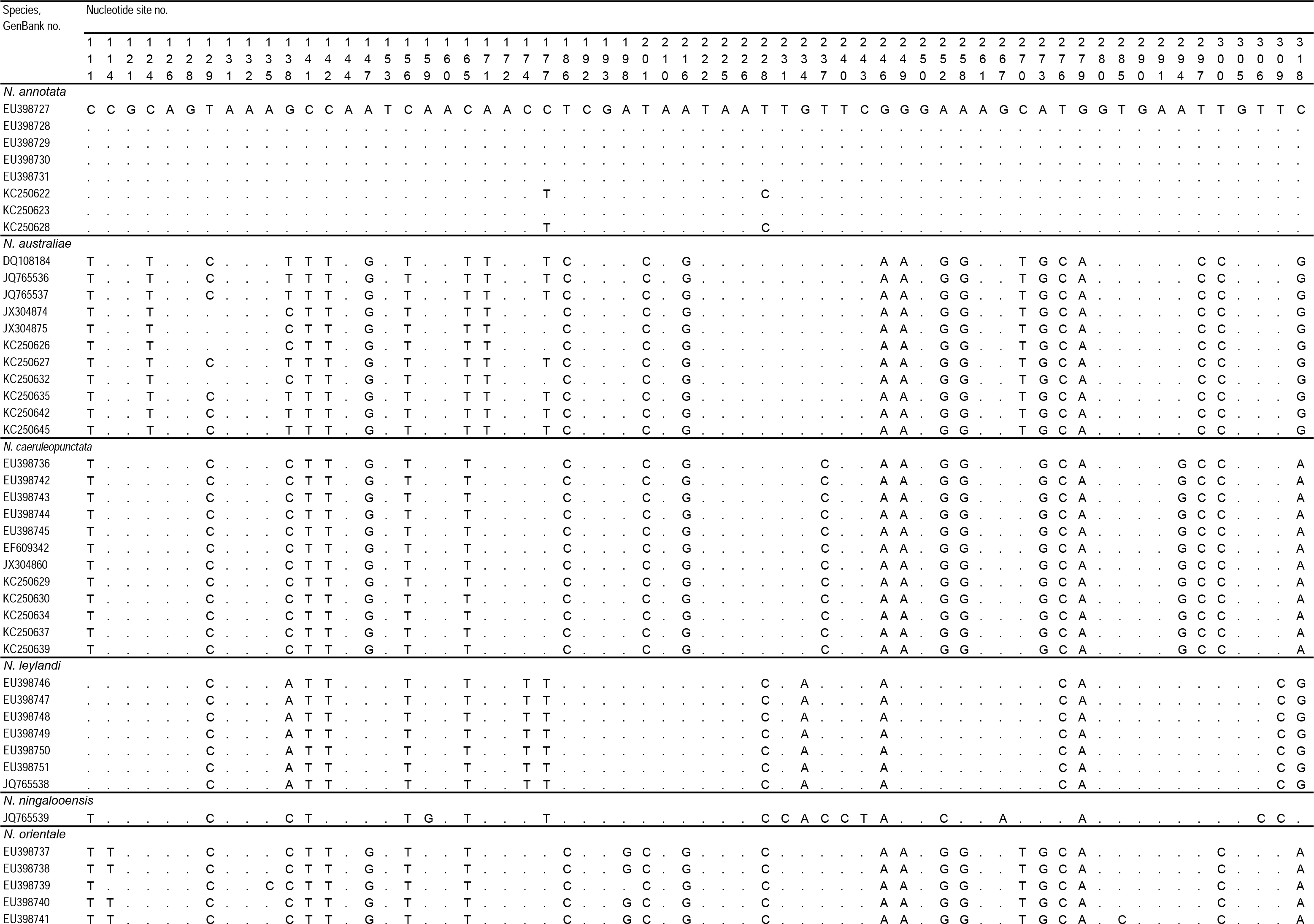
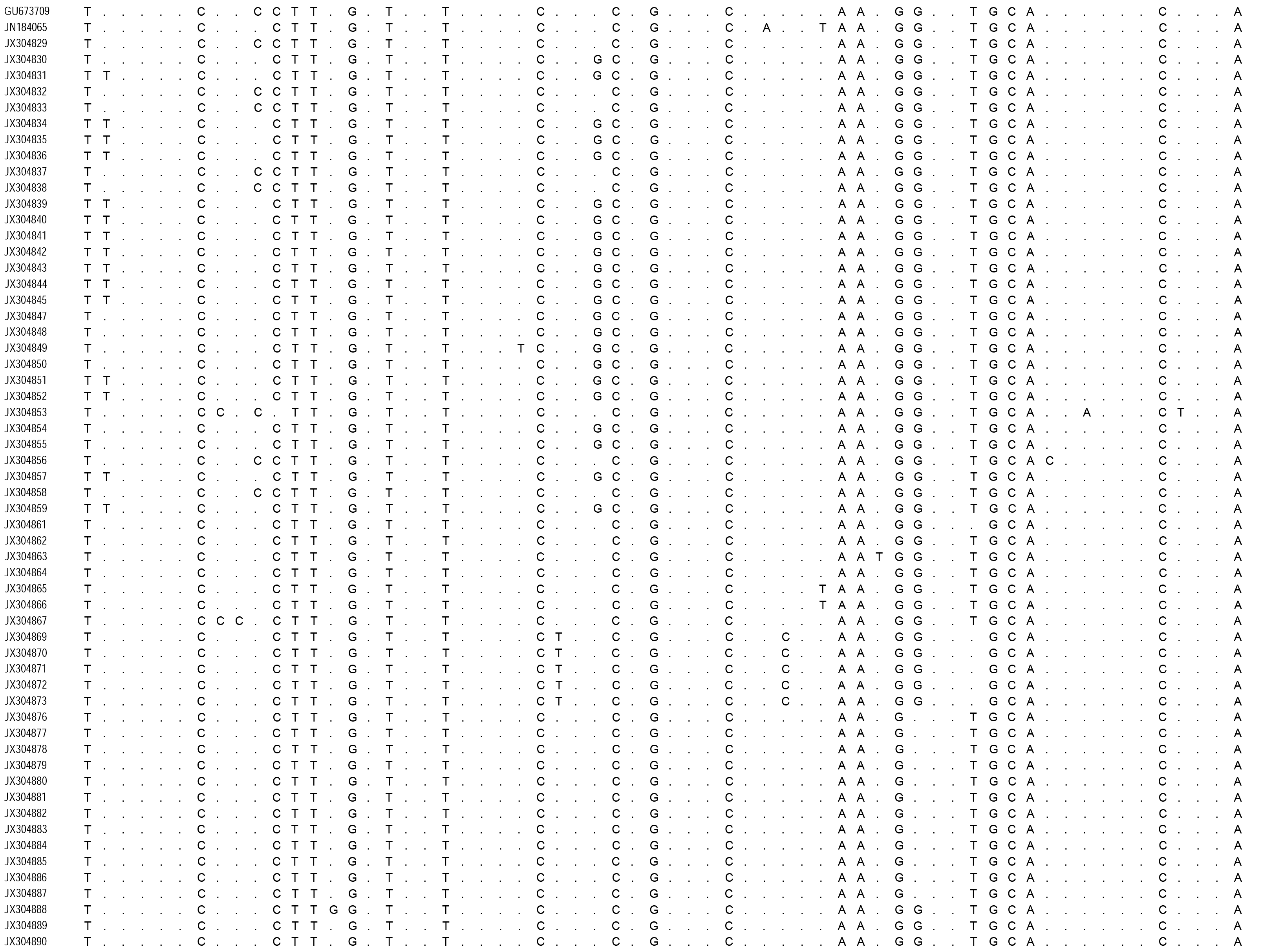
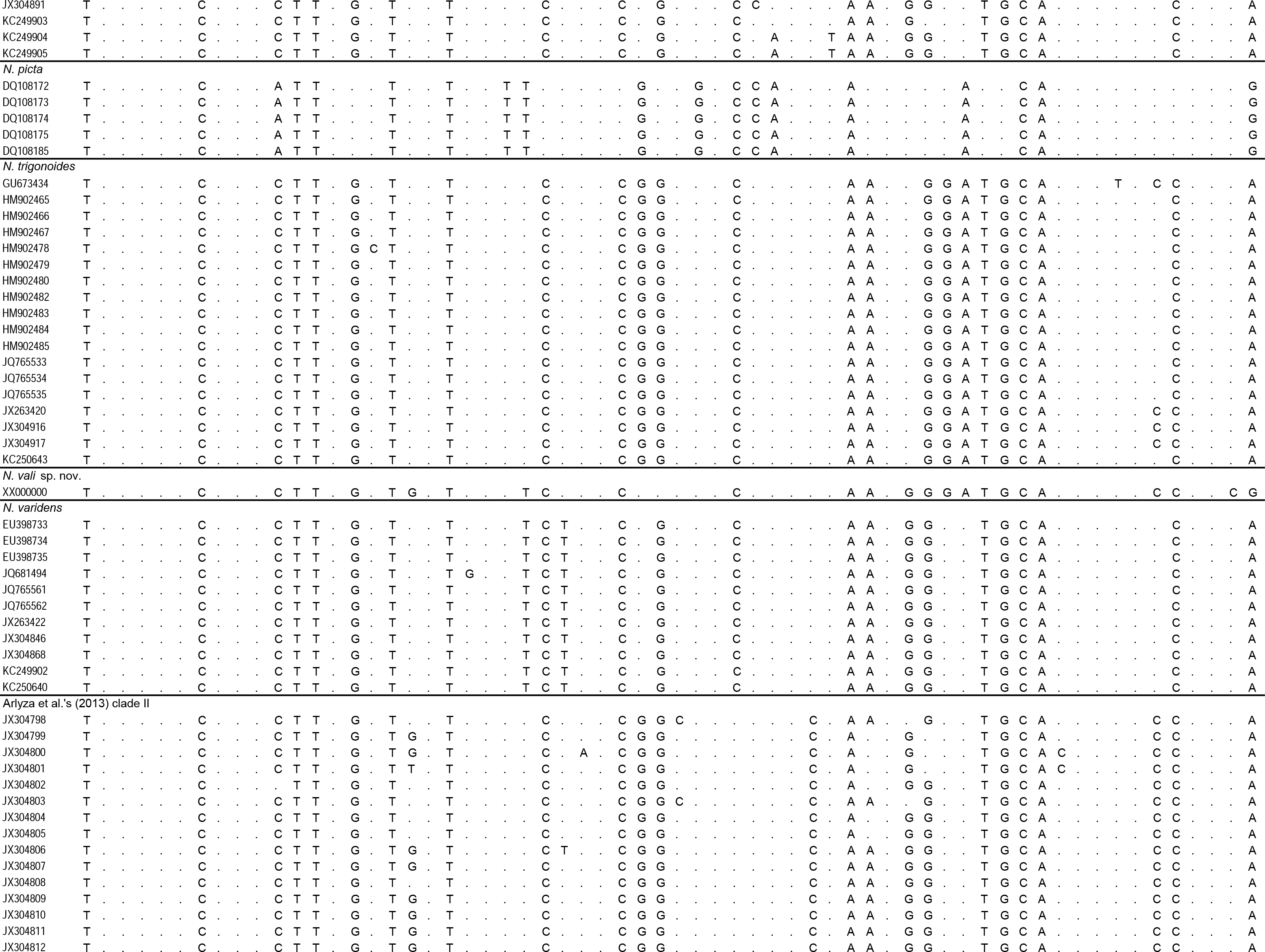
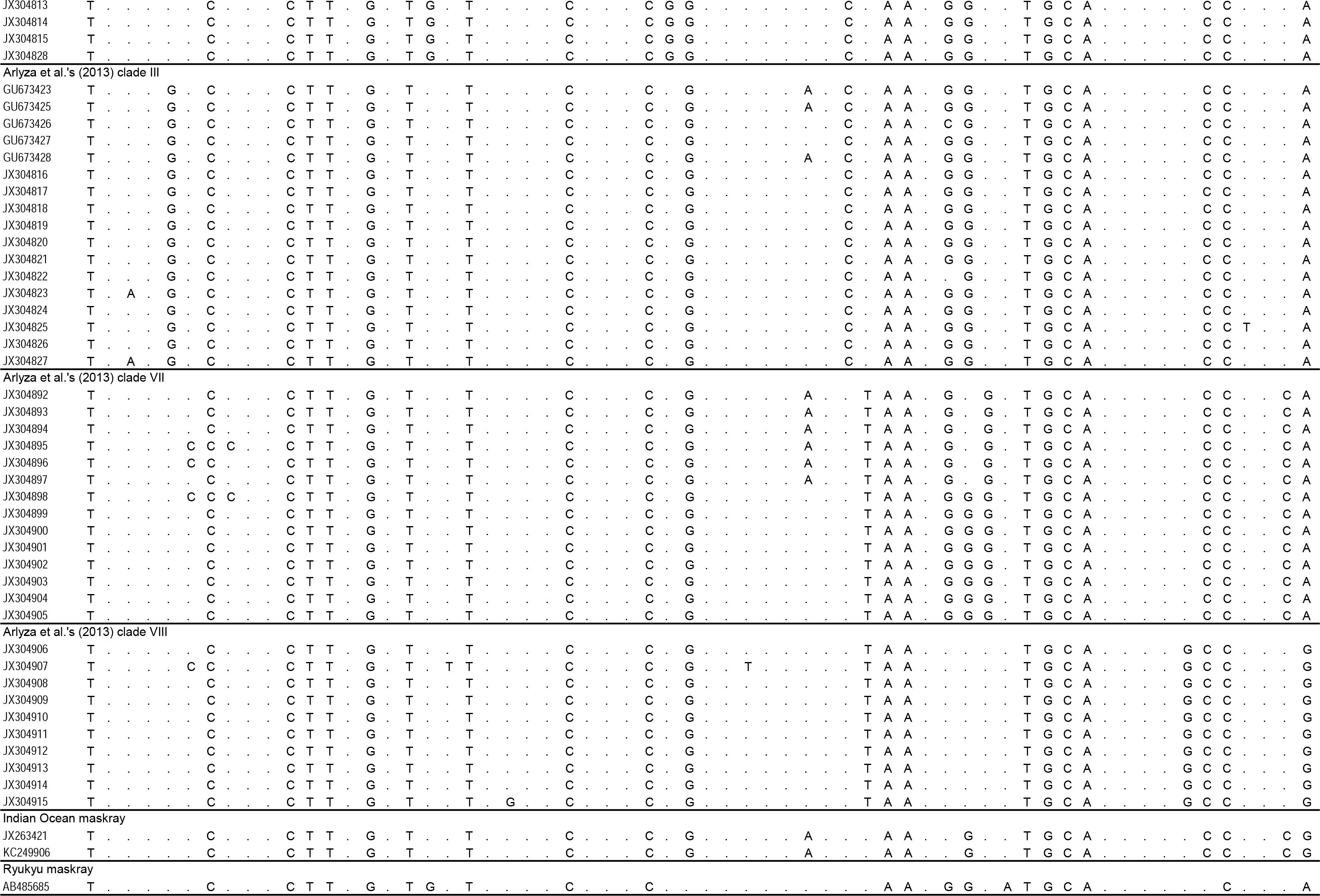
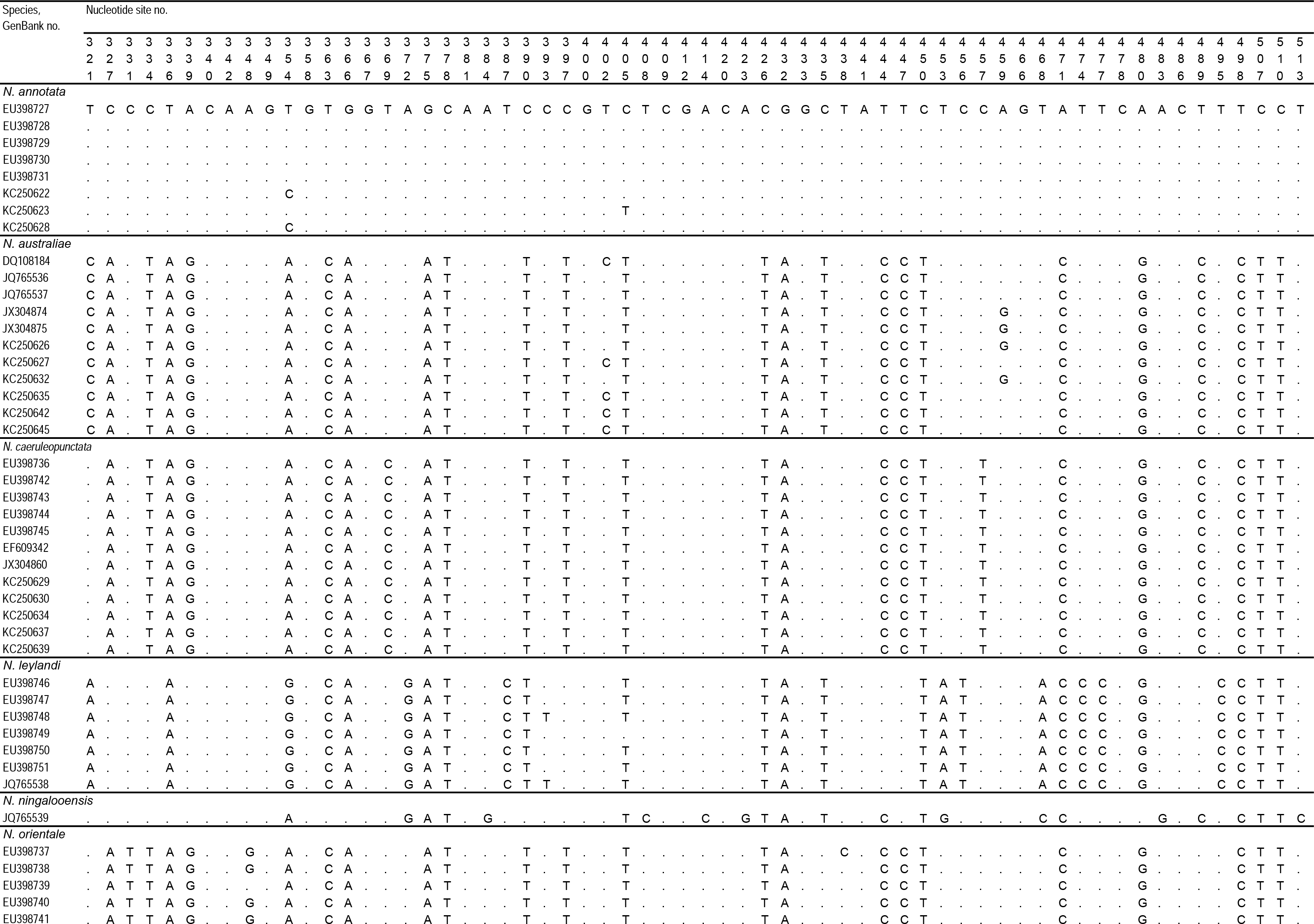
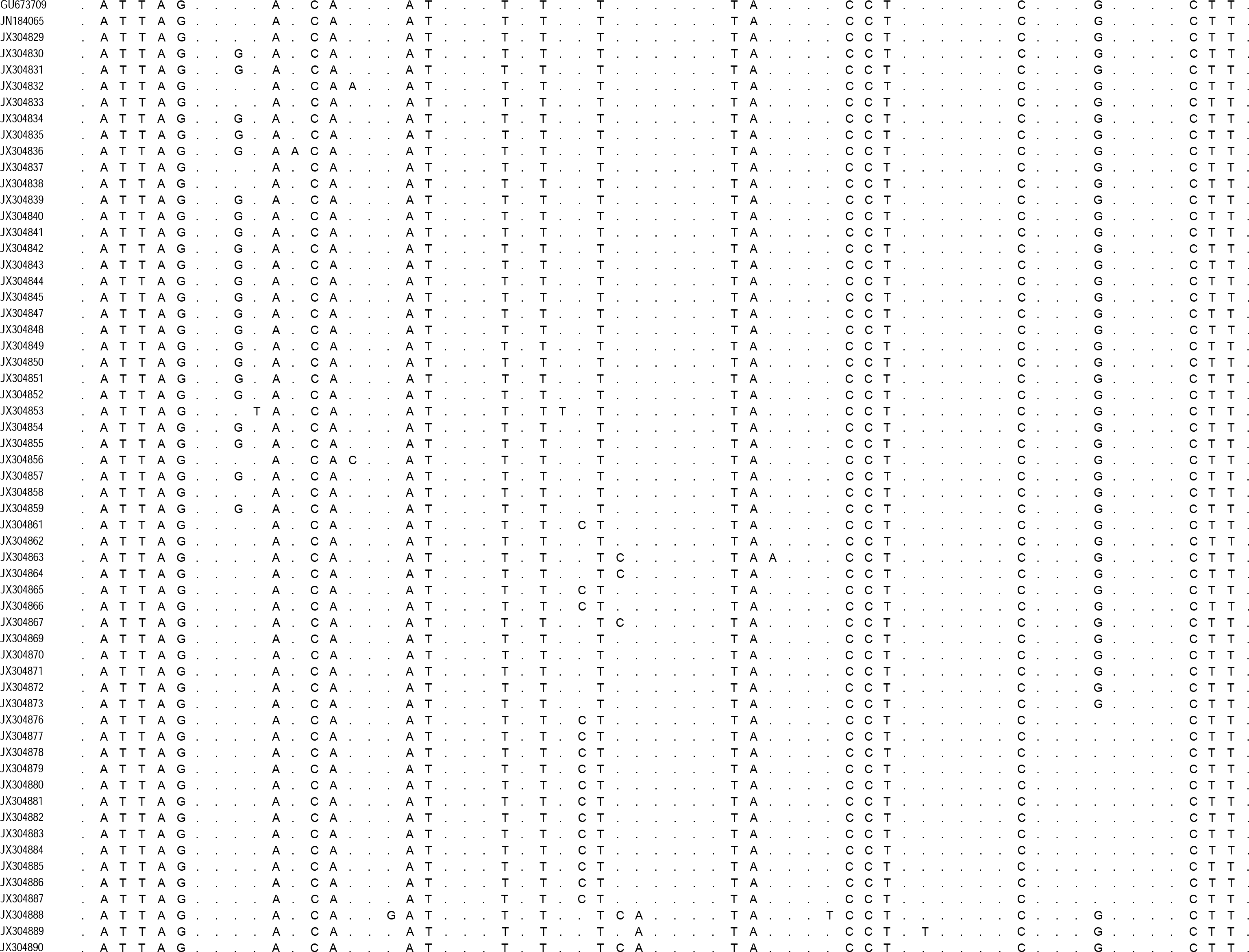
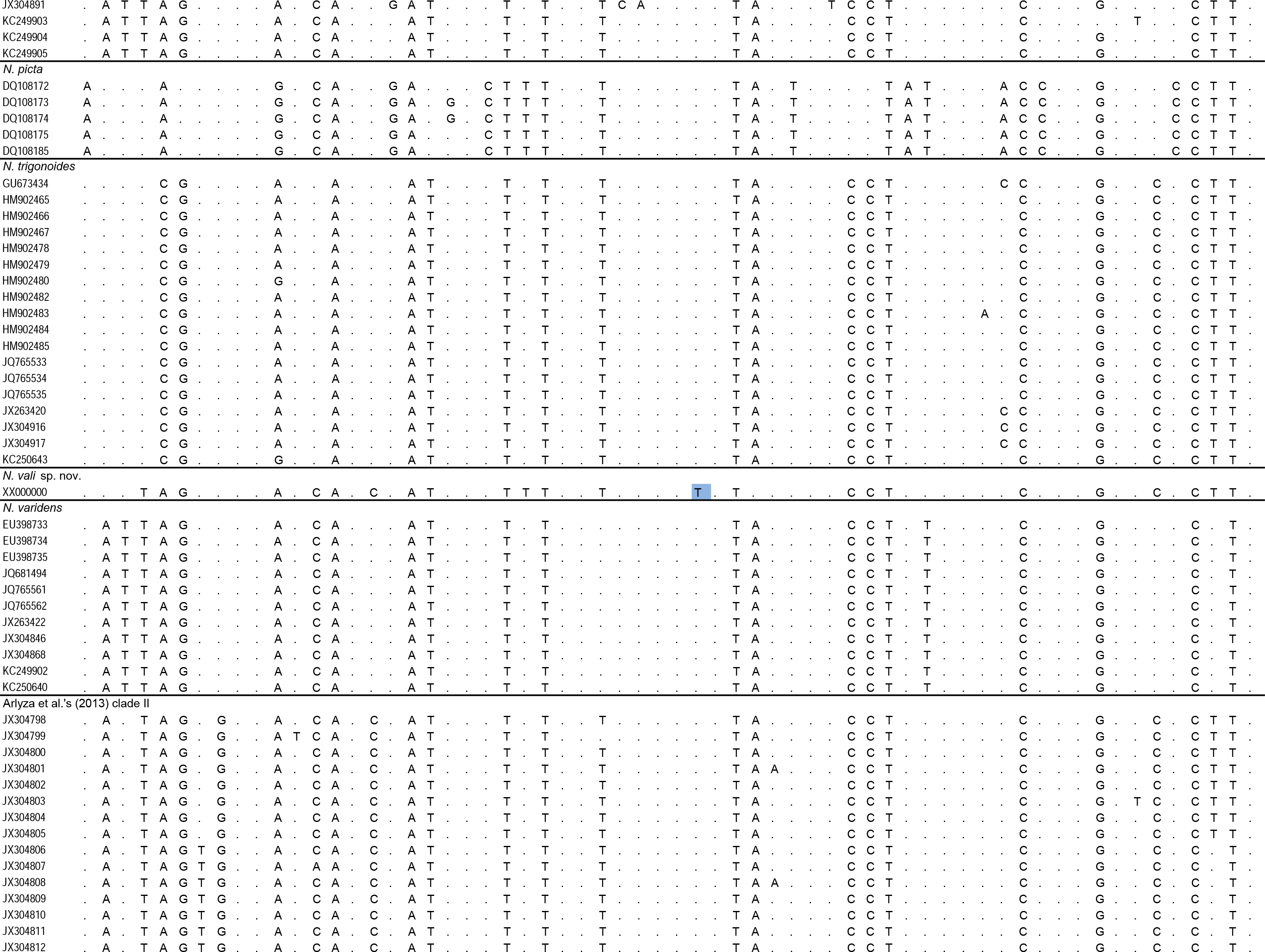
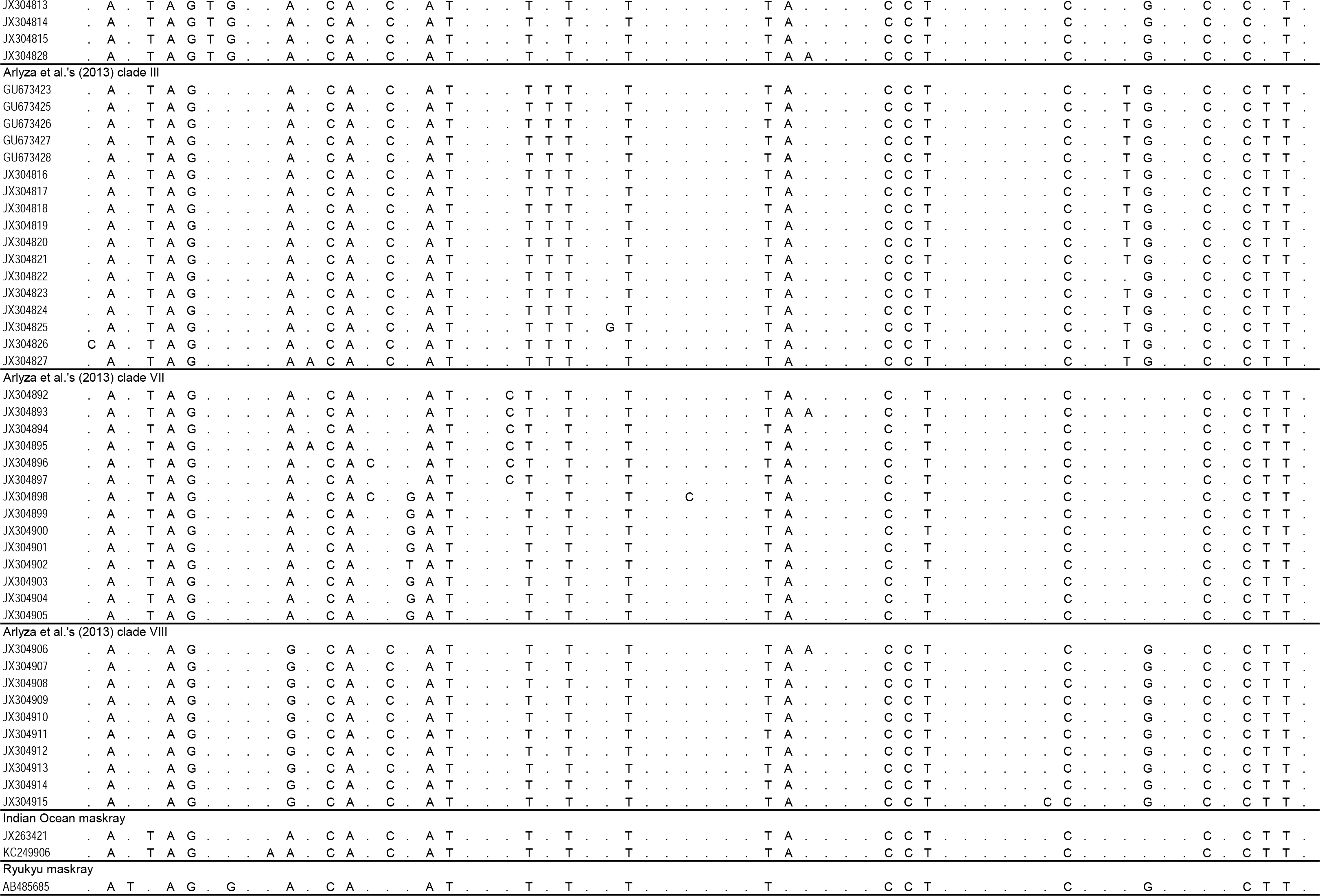
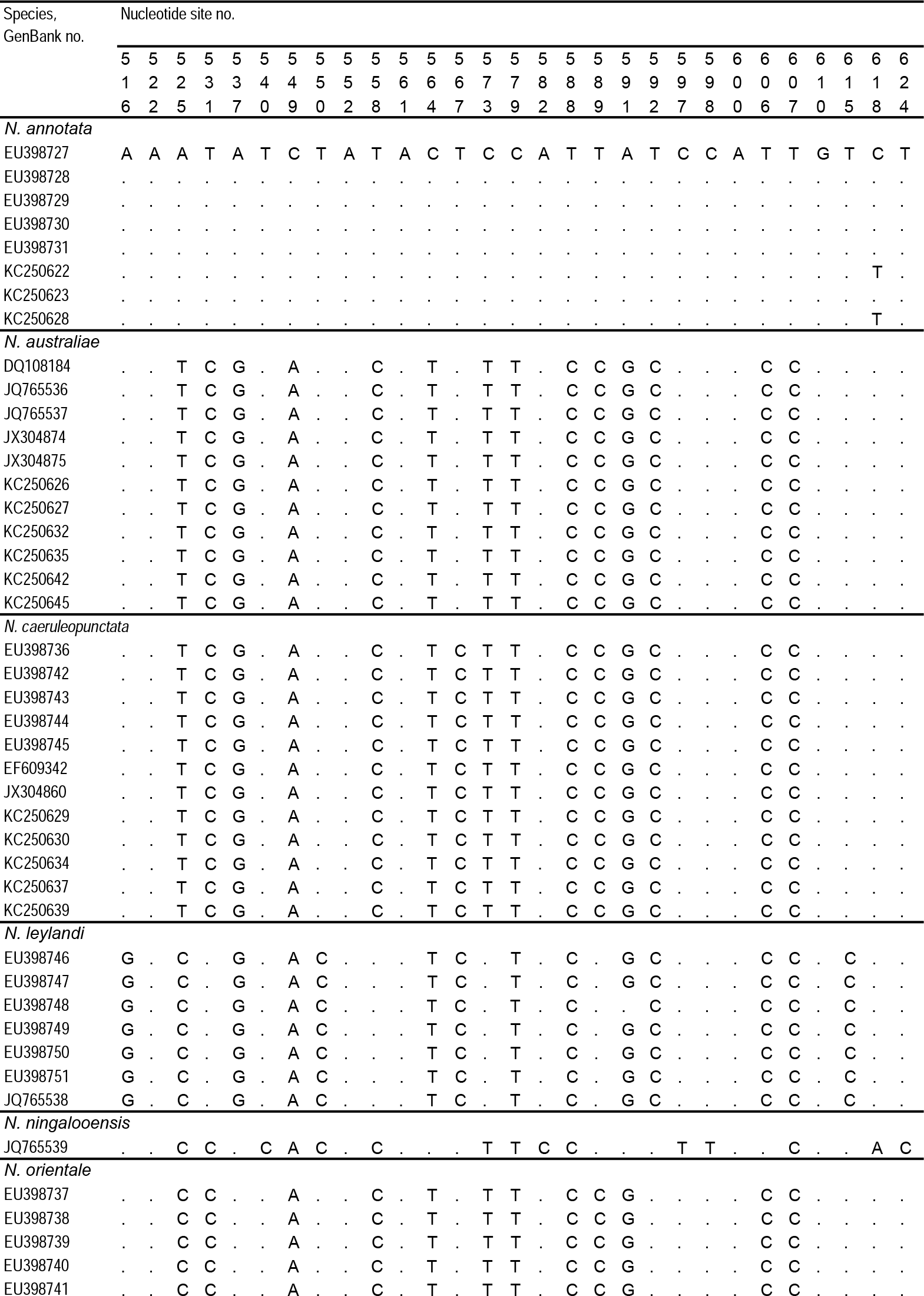
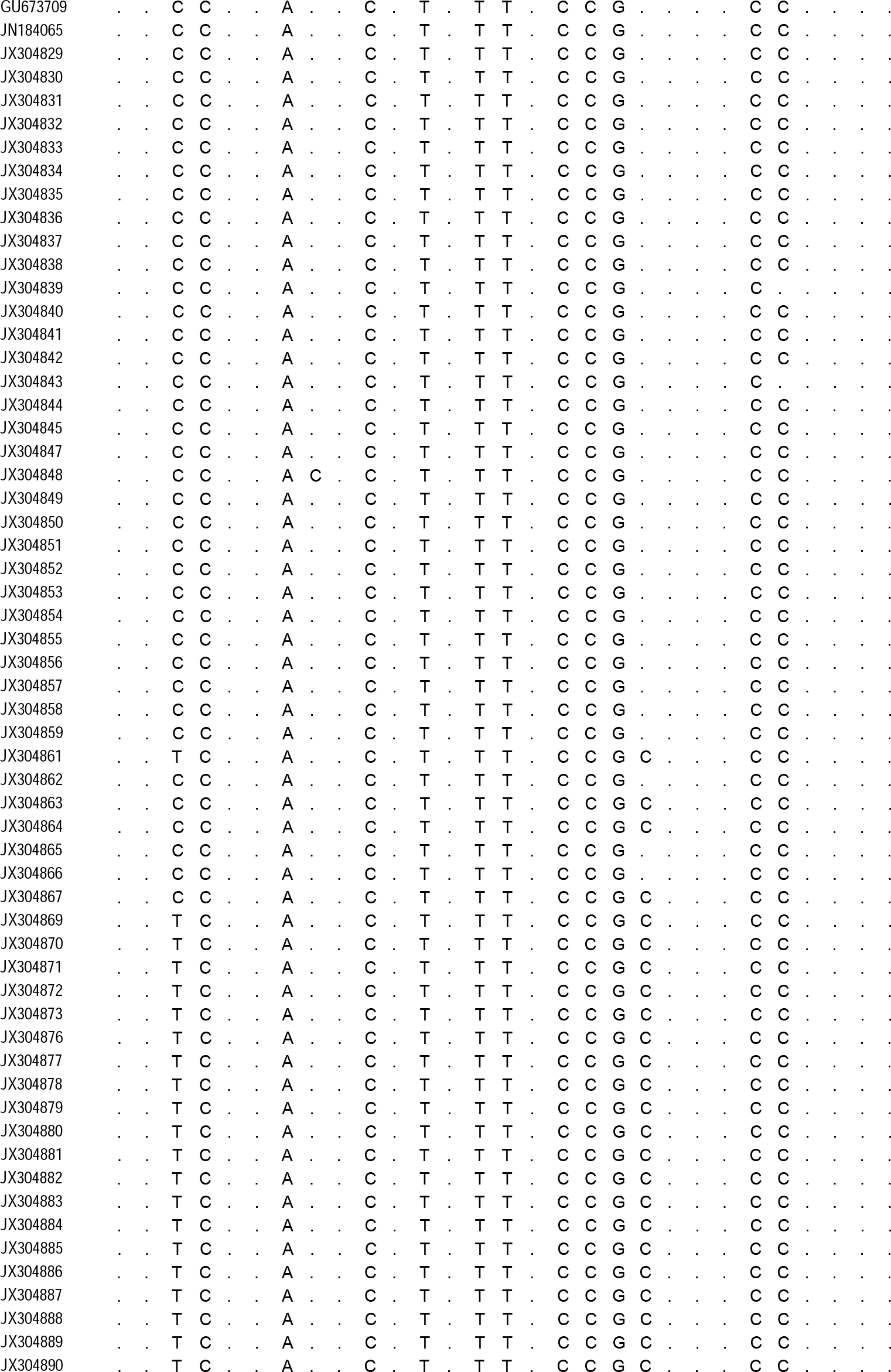
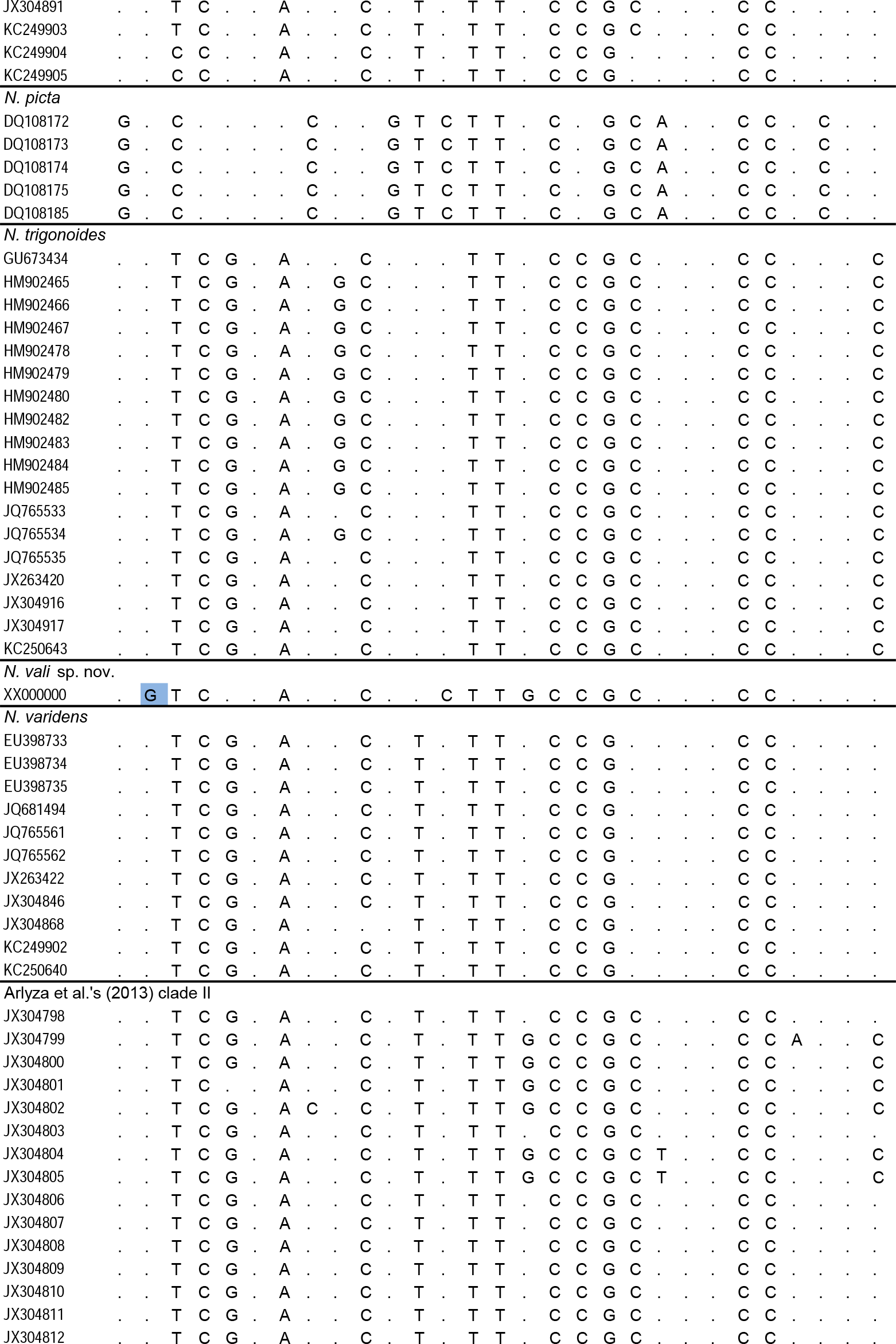
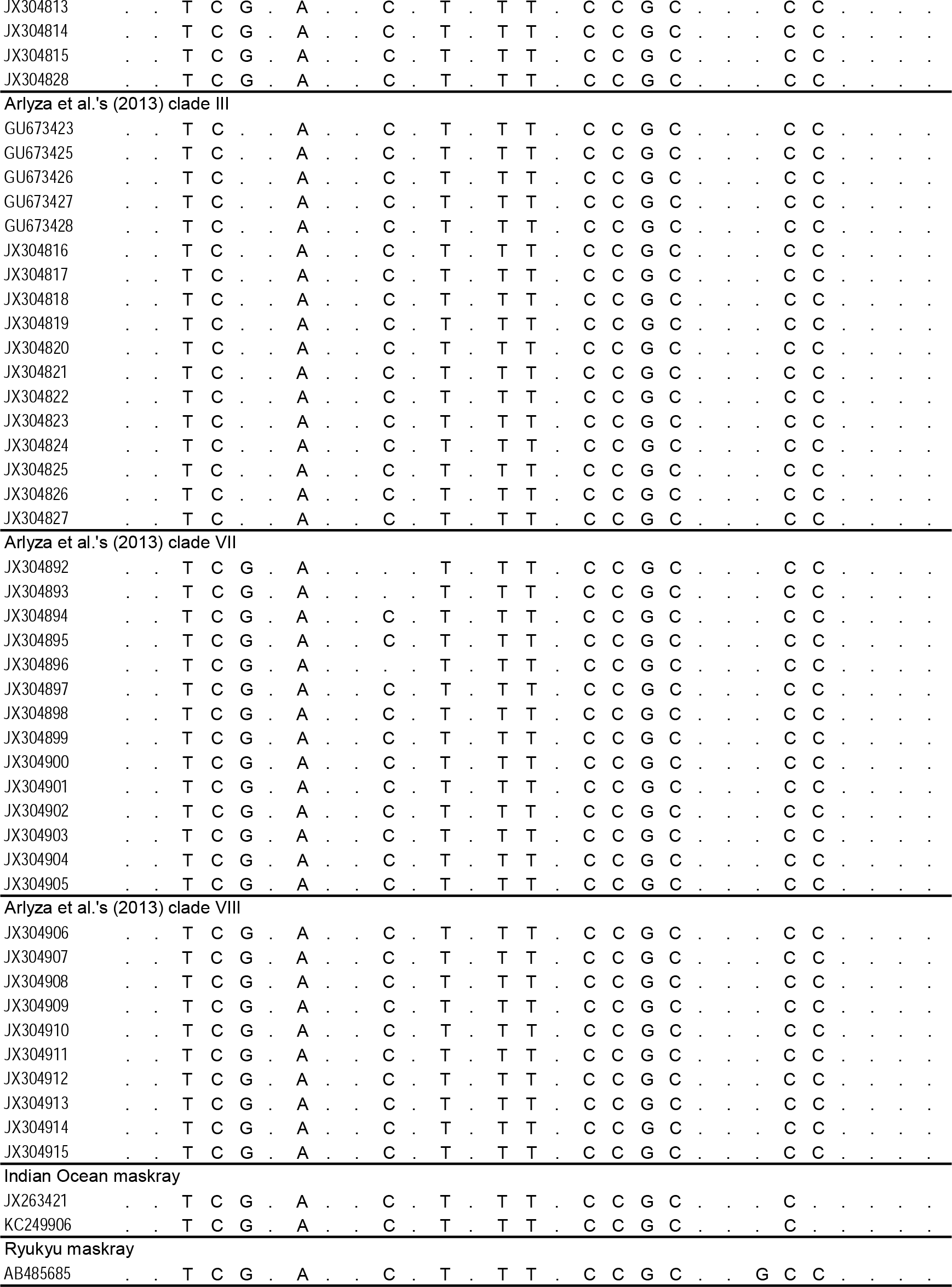
Variable nucleotide sites at the *CO1* locus that distinguish *Neotrygon vali* sp. nov. from congeneric species. Nucleotides diagnostic of *N. vali* sp. nov. are highlighted in blue. Nucleotide sites numerotated starting from the origin of the *CO1* gene in *N. orientale*, GenBank accession no. JN184065. The fragment used in this alignment is 519 bp long, spanning nucleotide sites 106-624.

**Supplementary Table S2.**
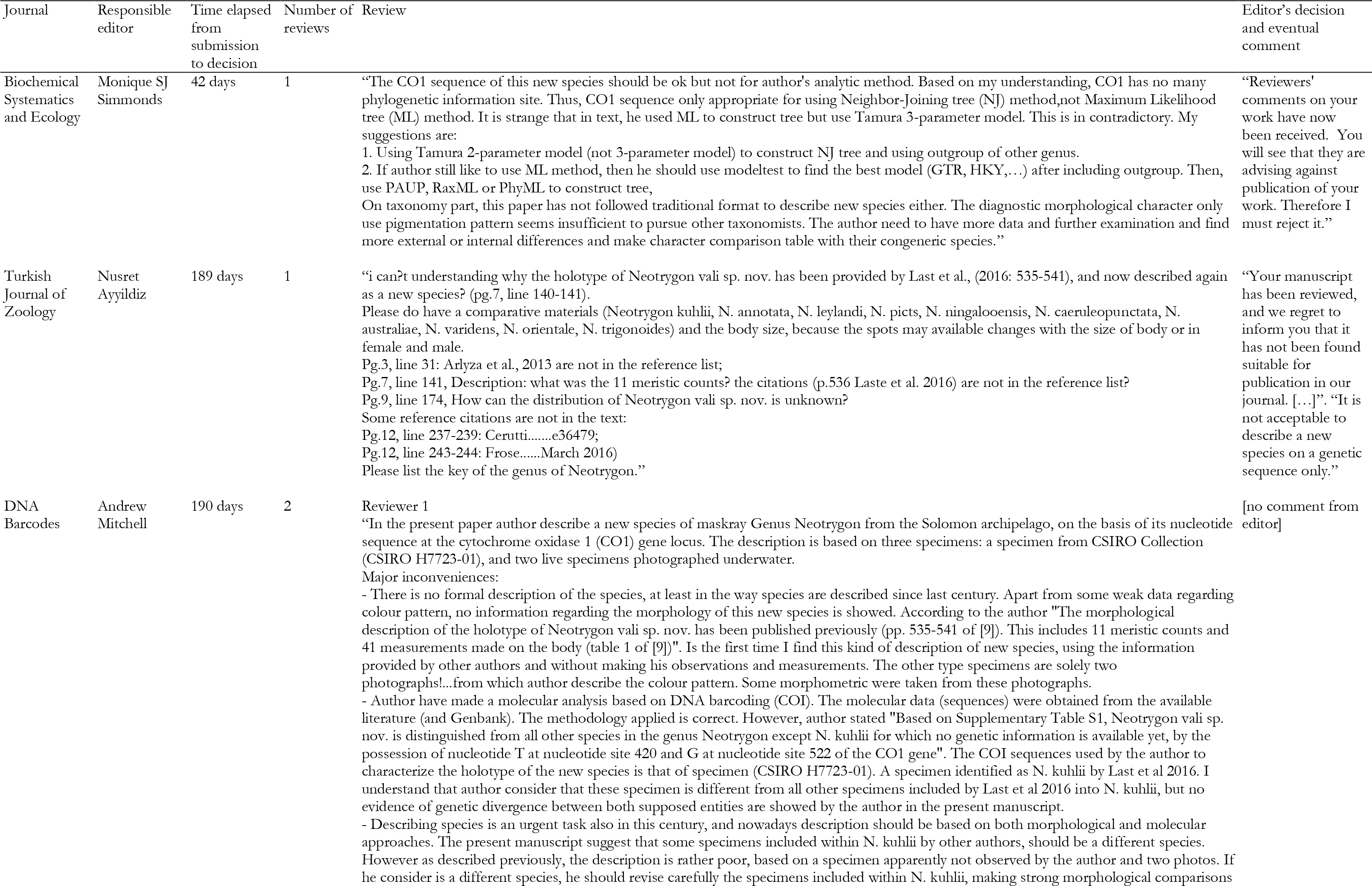
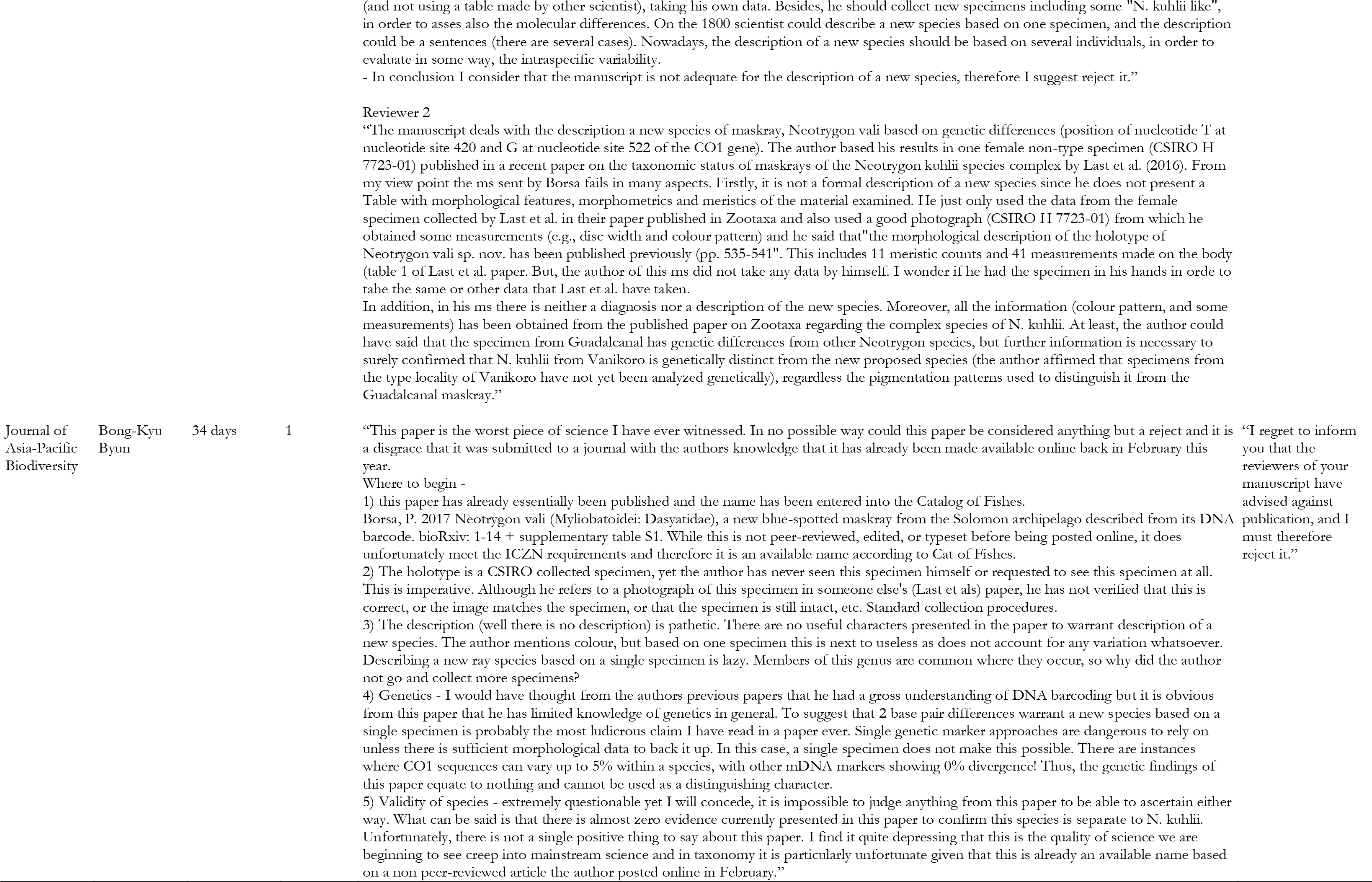
Reviews received by this manuscript, from four peer-reviewed journals to which it has been successively submitted, and each time rejected

## REFERENCES

Arlyza IS, Shen K-N, Durand J-D, Borsa P. 2013a. Mitochondrial haplotypes indicate parapatric-like phylogeographic structure in blue-spotted maskray (*Neotrygon kuhlii*) from the Coral Triangle region. J. Hered. 104:725–733.

Arlyza IS, Shen K-N, Solihin DD, Soedharma D, Berrebi P, Borsa P. 2013b. Species boundaries in the *Himantura uarnak* species complex (Myliobatiformes: Dasyatidae). Mol. Phyl. Evol. 66:429–435.

Aschliman NC, Nishida M, Miya M, Inoue JG, Rosana KM, Naylor GJP. 2012. Body plan convergence in the evolution of skates and rays (Chondrichthyes: Batoidea). Mol. Phyl. Evol. 63:28–42.

Borsa P. 2017. Comments on "Annotated checklist of the living sharks, batoids and chimaeras (Chondrichthyes) of the world, with a focus on biogeographical diversity" (Weigmann, 2016). J. of Fish Biol. 90:1170–1175.

Borsa P, Arlyza IS, Chen W-J, Durand J-D, Meekan MG, Shen K-N. 2013a. Resurrection of New Caledonian maskray *Neotrygon trigonoides* (Myliobatoidei: Dasyatidae) from synonymy with *N. kuhlii*, based on cytochrome-oxidase I gene sequences and spotting patterns. C. R. Biol. 336:221–232.

Borsa P, Arlyza IS, Hoareau TB, Shen KN. In press. Diagnostic description and geographic distribution of four new cryptic species of the blue-spotted maskray species complex (Myliobatoidei: Dasyatidae; *Neotrygon* spp.) based on DNA sequences. J. Oceanol. Limnol.. doi: 10.1007/s00343-018-7056-2

Borsa P, Béarez P. 2016. Notes on the origin of Müller and Henle’s illustration and type material of the blue-spotted maskray *Neotrygon kuhlii* (Myliobatoidei: Dasyatidae). Cybium 40:255–258.

Borsa P, Durand J-D, Chen W-J, Hubert N, Muths D, Mou-Tham G, Kulbicki M. 2016a. Comparative phylogeography of the western Indian Ocean reef fauna. Acta Oecol. 72:72–86.

Borsa P, Durand J-D, Shen K-N, Arlyza IS, Solihin DD, Berrebi P. 2013b. *Himantura tutul* sp. nov. (Myliobatoidei: Dasyatidae), a new ocellated whipray from the tropical Indo-West Pacific, described from its cytochrome-oxidase I gene sequence. C. R. Biol. 336:82–92.

Borsa P, Shen K-N, Arlyza IS, Hoareau TB. 2016b. Multiple cryptic species in the blue-spotted maskray (Myliobatoidei: Dasyatidae: *Neotrygon* spp.): an update. C. R. Biol. 339:417–426.

Castelnau F de. 1873. Contribution to the ichthyology of Australia. Proc. Zool. Acclim. Soc. Vic. 2:37–158.

Eschmeyer WN, Fricke R, van der Laar R. 2016. Catalog of fishes: genera, species, references, electronic version. Available at: http://researcharchive.calacademy.org/research/ichthyology/catalog/ [Date accessed: 23 March 2016]

Froese R, Pauly D (editors). 2016. FishBase, World Wide Web electronic publication. Accessible at: http://www.fishbase.org/ [Date accessed: 12 March 2016]

Garman S. 1885. Notes and descriptions taken from selachians in the U. S. National Museum. Proc. U. S. Natl. Mus. 8:39–44.

Hall TA. 1999. BIOEDIT: a user-friendly biological sequence alignment editor and analysis program for Windows 95/98/NT. Nucl. Acids Symp. Ser. 41:95–98.

International Commission on Zoological Nomenclature. 2012. Amendment of Articles 8, 9, 10, 21 and 78 of the International Code of Zoological Nomenclature to expand and refine methods of publication. Bull. Zool. Nomencl. 69:161–169.

Jordan DS. 1888. A manual of the vertebrate animals of the northern United States, including the district north and east of the Ozark mountains, south of the Laurentian hills, north of the southern boundary of Virginia, and east of the Missouri river, inclusive of marine species. Fifth edition, entirely rewritten and much enlarged. Chicago: A. C. McClurg and company. 375 pp.

Last PR. 1987. New Australian fishes. Part 14. Two new species of *Dasyatis* (Dasyatididae). Mem. Natl. Mus. Vic. 48:57–61.

Last P, White WT, Puckridge M. 2010. *Neotrygon ningalooensis* n. sp. (Myliobatoidei: Dasyatidae), a new maskray from Australia. Aqua Int. J. Ichthyol. 16:37–50.

Last PR, White WT, Séret B. 2016. Taxonomic status of maskrays of the *Neotrygon kuhlii* species complex (Myliobatoidei: Dasyatidae) with the description of three new species from the Indo-West Pacific. Zootaxa 4083:533–561.

Müller J, Henle FGJ. 1841. Systematische Beschreibung der Plagiostomen, mit sechzig Steindrucktafeln. Berlin: Veit und Comp. xxii+200 pp, 60 pl.

Jörger KM, Schrödl M. 2013. How to describe a cryptic species? Practical challenges of molecular taxonomy. Front. Zool. 10:59.

Last PR, White WT. 2008. Resurrection of the genus *Neotrygon* Castelnau (Myliobatoidei: Dasyatidae) with the description of *Neotrygon picta* sp. nov., a new species from northern Australia. CSIRO Mar. Atm. Res. Pap. 22:315–325.

Naylor GJP, Caira JN, Jensen K, Rosana KAM, White WT, Last PR. 2012. A DNA sequence-based approach to the identification of shark and ray species and its implications for global elasmobranch diversity and parasitology. Bull. Am. Mus. Nat. Hist. 367:1–262.

Puckridge M, Last PR, White WT, Andreakis N. 2013. Phylogeography of the Indo-West Pacific maskrays (Dasyatidae, *Neotrygon*): a complex example of chondrichthyan radiation in the Cenozoic. Ecol. Evol. 3:217–232.

Randall JE. 2005. Reef and shore fishes of the South Pacific, New Caledonia to Tahiti and the Pitcairn Islands. Honolulu: University of Hawai’i Press. xii+707 pp.

Tamura K. 1992. Estimation of the number of nucleotide substitutions when there are strong transition-transversion and G + C-content biases. Mol. Biol. Evol. 9:678–687.

Tamura K, Stecher G, Peterson D, Filipski A, Kumar S. 2013. MEGA6: Molecular Evolutionary Genetics Analysis version 6.0. Mol. Biol. Evol. 30, 2725–2729

Truper HG, De’Clari L. 1997. Taxonomic note: necessary correction of specific epithets formed as substantives (nouns) “in apposition”. Int. J. Syst. Bacteriol. 47:908–909.

Ward RD, Holmes BH, White WT, Last PR. 2008. DNA barcoding Australasian chondrichthyans: results and potential uses in conservation. Mar. Freshw. Res. 59:57–71.

Yagishita N, Furumitsu K, Yamaguchi A. 2009. Molecular evidence for the taxonomic status of an undescribed species of *Dasyatis* (Chondrichthyes: Dasyatidae) from Japan. Species Divers. 14:157–164.

